# Increased heterogeneity and task-related reconfiguration of functional connectivity within a lexicosemantic network in autism

**DOI:** 10.1101/2021.11.22.469604

**Authors:** Apeksha Sridhar, R. Joanne Jao Keehn, Molly Wilkinson, Yangfeifei Gao, Michael Olson, Lisa E Mash, Kalekirstos Alemu, Ashley Manley, Ksenija Marinkovic, Annika Linke, Ralph-Axel Müller

## Abstract

Autism spectrum disorder (ASD) is highly heterogeneous in etiology and clinical presentation. Findings on intrinsic functional connectivity (FC) or task-induced FC in ASD have been inconsistent including both over- and underconnectivity and diverse regional patterns. As FC patterns change across different cognitive demands, a novel and more comprehensive approach to network architecture in ASD is to examine the *change* in FC patterns between rest and task states, referred to as reconfiguration. This approach is suitable for investigating inefficient network connectivity that may underlie impaired behavioral functioning in clinical disorders. We used functional magnetic resonance imaging (fMRI) to examine FC reconfiguration during lexical processing, which is often affected in ASD, with additional focus on interindividual variability. Thirty adolescents with ASD and a matched group of 23 typically developing (TD) participants completed a lexicosemantic decision task during fMRI, using multiecho-multiband pulse sequences with advanced BOLD signal sensitivity and artifact removal. Regions of interest (ROIs) were selected based on task-related activation across both groups, and FC and reconfiguration were compared between groups. The ASD group showed increased interindividual variability and overall greater reconfiguration than the TD group. An ASD subgroup with typical performance accuracy (at the level of TD participants) showed reduced similarity and typicality of FC during the task. In this ASD subgroup, greater FC reconfiguration was associated with increased language skills. Findings suggest that intrinsic functional networks in ASD may be inefficiently organized for lexicosemantic decisions and may require greater reconfiguration during task processing, with high performance levels in some individuals being achieved through idiosyncratic mechanisms.

**Highlights:**

FC reconfiguration is a comprehensive approach to examining network architecture
Functional networks are inefficiently organized for lexicosemantic decisions in ASD
ASD may require greater reconfiguration during task processing
Some ASD individuals achieve high performance through idiosyncratic mechanisms

## Introduction

Autism spectrum disorder (ASD) is a lifelong neurodevelopmental disorder characterized by impaired social interaction and communication, as well as restricted or repetitive patterns of behavior, interests, and activities (American Psychiatric Association, 2013). Neural network connectivity in ASD has commonly been studied using functional Magnetic Resonance Imaging (fMRI), which measures changes in the blood-oxygen-level-dependent (BOLD) signal (Greicius et al., 2003; Schipul et al., 2012). Functional connectivity (FC) is conventionally measured based on BOLD correlations between regions. Previous studies examining intrinsic (resting state) FC (Van Dijk et al., 2009) and task-induced FC in ASD have yielded inconsistent results (Hernandez et al., 2015).

Although there is overall consensus that FC in ASD is atypical, the direction of FC changes remains under debate (Mohammad-Rezazadeh et al., 2016). Some studies suggest underconnectivity (e.g., Cheng et al., 2017; Jao Keehn et al., 2021), some predominant overconnectivity (e.g., Cerliani et al., 2015; Supekar et al., 2013), some both under and overconnectivity (e.g., Lynch et al., 2013; Monk et al., 2009), and some have failed to detect differences (e.g., Nomi & Uddin, 2015; Tyszka et al., 2014). While some of these inconsistencies may reflect regional differences, methodological choices (e.g., testing intrinsic vs. versus task-induced FC) likely play a critical role (Jones et al., 2010; Müller et al., 2011). For example, Nair and colleagues (2014) found that underconnectivity findings in ASD tended to be associated with inclusion of task effects, but overconnectivity with intrinsic FC (resting or after statistical removal of task effects).

A novel approach to investigating network architecture examines the *change* in FC patterns across different cognitive states (rest and task-evoked conditions) (Hearne et al., 2017; Salehi et al., 2020). It is generally understood in the fMRI literature that FC patterns are state-dependent, changing with time and across different mental states (e.g., Gonzalez-Castillo et al., 2015; Telesford et al., 2016). Some recent studies have examined the degree of FC changes between rest and task, referred to as FC *reconfiguration* (e.g., Hearne et al., 2017; Salehi et al., 2020). Reconfiguration, which reflects changes in resting state FC (intrinsic network architecture) that occur during domain-specific processing, provides a more comprehensive view of network connectivity. Thus, low reconfiguration may indicate that intrinsic architecture easily adapts to task processing without major ‘neural effort’ (Hearne et al., 2017), whereas high reconfiguration may reflect major changes in FC due to inefficient intrinsic network organization. A method focusing on FC reconfiguration can therefore serve to investigate non-optimized network connectivity that may underlie impaired behavioral functioning in clinical disorders such as ASD.

The TD brain is thought to require limited reconfiguration while performing tasks of moderate difficulty, due to generally efficient intrinsic architecture (Hearne et al., 2017). In addition, the level of task-related network reconfiguration in TD children has been found to be negatively associated with cognitive performance (with greater FC reconfiguration linked to poorer performance) in several domains, including working memory (Braun et al., 2015; Vatansever et al., 2015, 2017), attention (Shine et al., 2016), cognitive control (Dwyer et al., 2014), and general intelligence (Schultz & Cole, 2016). Hearne and colleagues (2017) have further suggested that reconfiguration increases when the system is pushed to the limits. These limits may be lower in ASD, with greater reconfiguration required for high (or neurotypical) levels of task performance. For example, Uddin and colleagues (2015) found that low FC reconfiguration in children with ASD was associated with severity of restricted and repetitive behaviors, presumably due to behavioral inflexibility. Here, we aimed to extend these findings to a different domain by employing a lexical decision task that involved additional processes including cognitive flexibility and executive functioning, which are often impaired in ASD (e.g., Van Eylen et al., 2011; Verhoeven et al., 2010). Previous behavioral studies have reported atypical performance on lexicosemantic decision and executive tasks in ASD (e.g., de Vries & Geurts, 2012; Ellis Weismer et al., 2018; Kamio et al., 2007), but the underlying neural network connectivity remains poorly understood.

Inconsistent findings of network connectivity in ASD may also be in part explained by high levels of heterogeneity (and cohort effects in limited samples). It has been proposed that the ASD brain may be characterized by variability of FC patterns across ASD individuals, referred to as ‘idiosyncrasy’ (Hahamy et al., 2015). FC patterns in adults with ASD have been found to be individually distinct or idiosyncratic during rest (Dickie et al., 2018; Hahamy et al., 2015; Hasson et al., 2009; Nunes et al., 2019). Such inter-individual variability may relate to findings of increased *intra*-individual variability in evoked cortical responses and spatio-temporal responses in ASD (Byrge et al., 2015; Dinstein et al., 2012).

In the current study, we used fMRI to examine FC reconfiguration associated with lexical processing in adolescents with ASD compared to a matched TD group. In view of evidence of high levels of heterogeneity in ASD, we also tested interindividual variability across task-dependent FC and resting state FC. We examined whether ASD participants who were able to perform at neurotypical levels differed from those whose performance was distinctly below neurotypical levels. We hypothesized that the ASD group would show greater reconfiguration and greater heterogeneity of resting state FC, task-induced FC, and reconfiguration compared to the TD group. Moreover, we predicted that the relation of FC reconfiguration with task performance, executive functioning, and language ability would be negative in the TD group, but positive in the ASD group.

## Methods

### Participants

The current study included 30 adolescents with ASD and 23 TD peers. Groups did not differ on gender, age, handedness, or non-verbal IQ (Table 1). Participants with ASD met diagnostic criteria according to the DSM-5 (American Psychiatric Association, 2013), the Autism Diagnostic Interview-Revised (ADI-R; Lord et al., 1994), the Autism Diagnostic Observation Schedule, 2nd edition (ADOS-2; Lord et al., 2012), and expert clinical decision. Participants diagnosed with any neurological disorder other than ASD (e.g., seizures, fragile X) or other comorbid disorders (e.g., Tourette’s syndrome) were excluded from the study. One ASD participant with co-occurring depression was not excluded due to the high prevalence of such conditions in autism (DeFilippis, 2018). All participants reported their primary spoken language as English, and participants with reported primary spoken language other than English before age 5 years were excluded to minimize confounds related to bilingualism (Gasquoine, 2016).

**Table 1.** Participant Demographics

| | ASD (n = 30) | | TD (n =23) | | $\chi^2$ | *p*-values | % Difference |
| --- | --- | --- | --- | --- | --- | --- | --- |
| **Gender** | 9 Female |  | 6 Female |  | 0.56 | 0.75 | 3.10 |
| **Handedness** | 1 left |  | 2 left |  | 1.82 | 0.40 | 5.10 |
|  | Mean (SD) | Range | Mean (SD) | Range | *t* (df) |  |  |
| **Age in years** | 15.3 (2.3) | 10.1 - 20.0 | 14.9 (1.9) | 12.1 -21.1 | 0.58 (51) | 0.57 | 3.10 |
| **RMSD Rest** | 0.07 (0.03) | 0.01- 0.13 | 0.06 (0.03) | 0.02 - 0.12 | 0.82 (51) | 0.42 | 8.72 |
| **RMSD Task** | 0.07 (0.02) | 0.03 - 0.13 | 0.06 (0.02) | 0.03 - 0.13 | 2.26 (51) | 0.03 | 21.91 |
| **WASI-II** |  |  |  |  |  |  |  |
| Nonverbal IQ | 110.3 (21.7) | 62-156 | 110.5 (11.9) | 80 -128 | -0.04 (51) | 0.97 | 0.20 |
| Verbal IQ | 106.1 (16.8) | 68 -134 | 110.8 (13.3) | 85 -135 | -1.10 (51) | 0.28 | 4.32 |
| Full Scale | 108.4 (19.3) | 54 -141 | 112.6 (13.3) | 88 -135 | -0.88 (51) | 0.38 | 3.77 |
| **WIAT-III** | 103.5 (18.8) | 59 -133 | 110.3 (8.1) | 100 -129 | -1.64 (51) | 0.09 | 7.22 |
| **ADOS-2<sup>†</sup>** |  |  |  |  |  |  |  |
| Social Affect | 8.8 (3.0) | 3-14 | -- | -- |  |  |  |
| RRB | 2.4 (1.9) | 0-9 | -- | -- |  |  |  |
| Total | 10.8 (3.4) | 6-20 | -- | -- |  |  |  |
| **ADI-R** |  |  |  |  |  |  |  |
| Social Interaction | 17.6 (4.3) | 11-27 | -- | -- |  |  |  |
| Communication | 13.8 (3.9) | 8-21 | -- | -- |  |  |  |
| Repetitive Behavior | 5.5 (2.1) | 1-8 | -- | -- |  |  |  |
| **BRIEF-2 GEC** | 65.7 (7.4) | 49-84 | 46.4 (8.0) | 36-62 | 8.46 (44) | <0.001 | 27.88 |
| **CELF-5 WC** | 35.2 (4.2) | 27-40 | 36.0 (2.4) | 31 -39 | -1.51 (51) | 0.14 | 7.10 |

Participants who received a standard score < 80 (i.e., 2 standard deviations below the median [50^th^ percentile]), based on age 12 year reading norms on the Word Reading subtest of the Wechsler Individual Achievement Test, 3^rd^ Edition (WIAT-III; Wechsler, 2009), or who obtained <60% accuracy on a screening task (described below) were also excluded. Among the final sample (N=53), 11 ASD (and 0 TD) participants reported psychotropic medication use (Supplemental Table S1). These participants were not excluded due to high rates of medication use in ASD (Schubart et al., 2014). All participants provided written informed assent, and parents or guardians provided written informed consent. The study was approved by the

### Neuropsychological Measures

Participants were administered a battery of age-appropriate neuropsychological tests, which assessed major areas of neuropsychological functioning including cognitive, perceptual, and social domains. At the initial appointment, participants completed the WIAT-III (Wechsler, 2009), the Wechsler Abbreviated Scale of Intelligence, 2^nd^ Edition (WASI II; Wechsler, 1999), and the Clinical Evaluation of Language Fundamentals, 5^th^ Edition (CELF-5; Wiig et al., 2013), among others. Parents or guardians completed questionnaires regarding the participant’s behavior and executive functioning such as the Behavior Rating Inventory of Executive Function, 2^nd^ Edition (BRIEF-2; Gioia et al., 2015).

### Experimental Paradigm

For the experimental task, participants were asked to distinguish between ‘animal words’ (AW; e.g., “cat”), ‘standard words’ (SW; i.e., moderately high frequency nouns from any semantic category other than animals; e.g., “chair”), and ‘pseudowords’ (PW; orthographically and phonologically legal letter strings without semantic content; e.g., “blont”). Accuracy and reaction time (RT) were recorded through button press responses (see below). Each 2 second trial included a stimulus presented for 500 ms followed by a 1500 ms fixation string (“xxxxxx”) to allow for response. One-second null trials (124 per run) consisting of the fixation string were also included. Standard words and animal words did not differ on age of acquisition (Kuperman et al., 2012). Additionally, conditions did not differ in number of letters or syllables (Supplemental Table S2).

**Table 2.** Task Performance per Sample and Between Sample Comparisons

|  |  | ASD |  | TD |  | *t* (df), *p*-value |
| --- | --- | --- | --- | --- | --- | --- |
|  |  | Mean (SE) | Range | Mean (SE) | Range |  |
| SW | Accuracy | 0.90 (0.02) | 0.67 – 1.0 | 0.97 (0.01) | 0.90 – 0.99 | -3.1(51), ** *p* = 0.003 |
|  | RT | 946.4 (25.7) | 630.2 – 1227.4 | 837.9 (28.7) | 582.0 – 1106.0 | 2.1 (51), * *p* = 0.04 |
| AW | Accuracy | 0.80 (0.03) | 0.45 – 1.0 | 0.90 (0.01) | 0.77 – 0.97 | -2.5 (51), * *p* = 0.02 |
|  | RT | 938.7 (20.9) | 679.0 – 1148.8 | 850.7 (26.6) | 601.3 – 1137.1 | 1.9 (51), *p* = 0.06 |
| PW | Accuracy | 0.91 (0.02) | 0.67 – 1.0 | 0.97 (0.02) | 0.68 – 1.0 | -1.6 (51), *p* = 0.11 |

Mean value, standard error (SE), range, *t*-statistic, degrees of freedom (df), and *p*-value of accuracy (calculated as number of correct responses divided by number of total responses) and response time (RT; in milliseconds) for each semantic category (SW, AW, PW) in each group (ASD, TD). \*\*\* *p*<.001, \*\* *p*<.01 \* *p*<.05.

The event-related fMRI design was created using the random stimulus function generator (RSFgen) in Analysis of Functional NeuroImages (AFNI Version 2.7.11; Cox, 1996; https://afni.nimh.nih.gov). Ten-thousand random permutations of the stimulus sequence were evaluated via 3dDeconvolve in AFNI, which can output the normalized standard deviation for each randomized sequence. The optimal sequence was selected with the lowest normalized mean standard deviation, a maximum of 5 sequential trials of any word category and a maximum of 3 sequential null trials. Two task runs with different stimuli were created using this optimal sequence. Participants were administered the same trial sequence except for 2 TD and 4 ASD individuals who required repeat scans and received an alternate sequence (to limit practice effects).

Prior to the actual MRI session, participants were familiarized with the task and scanner. They were first given instructions on how to respond to the stimuli and then completed a practice session of the task (36 trials total: 12 SW, 12 AW, 12 PW) on a laptop computer (Dell Precision M2800). Participants responded to SW by using their left index finger and AW using their left middle finger on different keys on the keyboard. They received feedback from the examiner immediately after each word stimulus to ensure they understood the instructions. During this appointment, participants were also familiarized with the MRI environment using a mock scanner to become accustomed to lying still inside a scanner. A second practice test without direct feedback was administered inside the mock scanner (90 trials total: 54 SW, 18 AW, 18 PW) and participants responded using their left hand on a two-button response pad (Fibre Optic Response Device). Those who scored below 60% accuracy were excluded from the study. The mock session consisted of word stimuli that differed from those presented during the functional MRI task.

At the start of the MRI session, participants were reminded of the task instructions and completed two more practice tasks on the same laptop computer, first with feedback, and then without feedback. Next, inside the scanner, a resting state scan was acquired during which participants were instructed to relax, stay awake, and keep their eyes centered on a white fixation cross presented on a black background using an LCD projector. In two following fMRI task runs, stimuli were presented (using Presentation software, v.22.1; Neurobehavioral Systems) on a 120 x 90 cm screen in front of the scanner, viewed through a front-facing mirror. Participants were instructed to respond to SW (90 trials per run) using their left index finger on the two-button response pad, to AW (30 trials per run) using their left middle finger, and to inhibit responses to words they had never seen before (i.e., PW; 30 trials per run). Participants were monitored with an in-bore camera during the experiment to ensure vigilance and continuous eyes-open status throughout resting and task scans.

### MRI Parameters

MRI scans were performed on a General Electric (GE) Discovery MR750 3.0T (GE Healthcare, Chicago) whole-body scanner with a 32-channel head coil at the University of California San Diego Center for Functional MRI. To minimize motion artifacts, combinations of foam pads for different head sizes were used. High-resolution structural images were acquired with a fast spoiled gradient echo (FSPGR) T1-weighted sequence (TR= 8.136ms, TE= 3.172 ms, flip angle= 8°; FOV = 25.6 cm, matrix = 256 x 192, voxel size = 1mm^3^, 172 slices). An accelerated multi-echo simultaneous multi-slice (MESMS) echo planar imaging (EPI) sequence (Cohen et al., 2020; Kundu et al., 2012; Olafsson et al., 2015) was used to acquire one resting state fMRI scan (309 volumes, 6:26 minutes) and two task runs (340 volumes, 7 minutes) with the following parameters: TR=1250ms; TEs=13.2, 30.3, 47.4ms; flip angle=60°; FOV=21.6cm; acquisition matrix=72x36; in-plane acceleration factor = 2; multiband acceleration factor = 3; 54 slices; voxel size=3mm^3^. The functional protocol slightly differed for 9 TD and 2 ASD participants (TR = 1100 ms; 45 slices; 340 volumes, 6 minutes 14 seconds for resting state; 386 volumes for task runs; all other parameters identical). To allow for magnetization to reach equilibrium, the first 9 time points of each run were discarded. Multiecho fMRI is not yet commonly used, but is increasingly recognized for its improved BOLD signal sensitivity and artifact removal while also allowing for high temporal resolution, compared with conventional single echo fMRI (Lynch et al., 2020). Notably, very few ASD studies have used a combined multiecho-multiband EPI sequence (King et al., 2018; Linke et al., 2020).

### fMRI Pre-processing

Functional images were processed using AFNI and FSL (v5.0; Smith et al., 2004), and filtered using MATLAB 2018a (The MathWorks, Inc.). To minimize susceptibility-induced distortions, two spin-echo EPI acquisitions with opposite phase encoding directions were used with FSL’s TOPUP tools (Smith et al. 2004). Rigid-body realignment was implemented using AFNI by registering each functional volume to the middle time point of the scan to adjust for motion. Functional data were then denoised using multi-echo independent component analysis to remove artifactual components (ME-ICA; Spreng et al., 2019; Kundu et al., 2013). As described by Olafsson and colleagues (2015), multi-echo weighted optimization and ME-ICA were performed using meica.py (<u>github.com/ME-ICA/me-ica</u>). EPIs from the three echoes were optimally combined (Kundu et al., 2013). Subsequently, functional images were co-registered to the anatomical scan via FSL’s FLIRT (Jenkinson and Smith, 2001) and standardized to the atlas space of the Montreal Neurological Institute (MNI) template using FSL’s nonlinear registration tool (FNIRT). The images were smoothed to a Gaussian full width at half-maximum (FWHM) of 6 mm via AFNI’s 3dBlurToFWHM. Lastly, resting fMRI data were filtered using a Butterworth bandpass filter (.008 < f < .08 Hz), while the task data were high-pass filtered (f > 0.01 Hz) to preserve any effects of the task that might also be observable at higher frequencies.

### Regions of Interest (ROIs)

Regions of interest (ROIs) were obtained using a one-sample linear contrast (SW+AW+PW>Null) across all participants from the fMRI task scans. A mixed-effects multilevel analysis (MEMA; Chen et al., 2012) was performed controlling for age, head motion, and task accuracy using the 3dMEMA function in AFNI. To control for false positive rates, randomization and permutation simulations were used to obtain cluster sizes using 3dttest++ in AFNI. All clusters at *p* < .001 (alpha = 0.05) were examined. Large clusters that included multiple brain regions were further thresholded to obtain smaller distinct brain regions. This resulted in 16 ROIs (minimum cluster size = 20 voxels; Figure 1; Supplemental Table S3).

**Figure 1.**
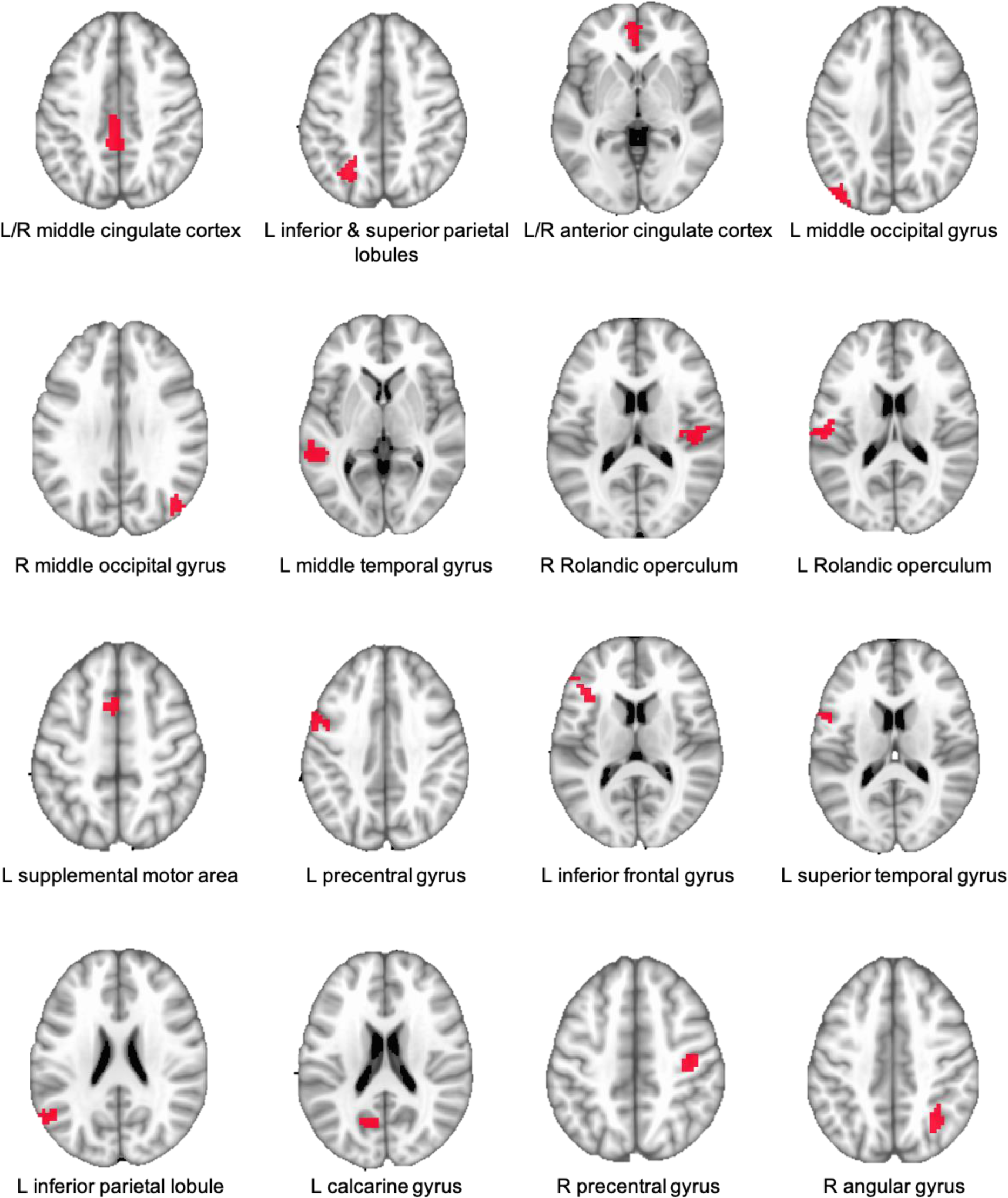
Regions of Interest (ROIs): L – left hemisphere; R – right hemisphere

**Table 3.** Task Performance and Comparisons Between Conditions per Group

|  |  | **ASD** | **TD** |
| --- | --- | --- | --- |
|  |  | *t* (df), *p*-value | *t* (df), *p*-value |
| **SW vs AW** | **Accuracy** | *t* (29) = 4.4, *** *p* < 0.001 | *t* (22) = 7.0, *** *p* < 0.001 |
|  | **RT** | *t* (29) = 0.8, *p* = 0.45 | *t* (22) = -1.0, *p* = 0.31 |
| **AW vs PW** | **Accuracy** | *t* (29) = -4.4, *** *p* < 0.001 | *t* (22) = -4.0, *** *p* < 0.001 |
| **PW vs SW** | **Accuracy** | *t* (29) = -0.3, *p* = 0.73 | *t* (22) = -0.9, *p* = 0.38 |

Paired samples *t*-test results of accuracy and response time (RT) between each semantic category (SW, AW, PW) for both groups (ASD, TD). \*\*\* *p* < .001, \*\* *p* < .01 \* *p* < .05.

### Subgroups

Groups differed significantly in task performance accuracy (*t* (51) = -2.72, *p* < 0.05), with some ASD participants performing at typical levels and others distinctly below the TD mean. Most TD participants performed above 90% accuracy. To better understand how potential group differences in FC and its reconfiguration are related to task performance, a cut-off of 90% accuracy was used to divide the ASD sample into those performing at typical levels (matched to the TD group in task accuracy) [typically-performing ASD subgroup: TP-ASD; n = 15, mean = 95%, standard deviation [std] = 0.03] and those with atypically low performance [low performing ASD subgroup: LP-ASD; n = 15, mean = 83%, std = 0.05]. Four TD participants with <90% accuracy (i.e., > 2 standard deviations below the TD group mean [0.94]) were excluded from the TD subgroup (TDs; n = 19, mean = 96%, std = 0.02) in all comparisons with ASD subgroups (refer to Supplemental Figure S1). Subgroups did not differ on age or non-verbal IQ (refer to Supplemental Tables S4 – S5).

### FMRI Functional Connectivity Analysis

Data analysis was conducted with MATLAB 2018B (The MathWorks, Inc.). From each ROI, a BOLD time series (averaged across all voxels within the ROI) was extracted. To obtain resting state FC and task FC, Pearson’s correlations were calculated for each participant between time series from all pairs of ROIs, for each of the two task runs and for the resting state scan.

Correlation coefficients were normalized using Fisher’s *z*-transformation. Since the two task runs did not significantly differ, FC (z’) from both runs was averaged into one task FC matrix.

### Functional Connectivity Reconfiguration Analysis

FC reconfiguration was calculated for each participant as the absolute difference between resting state FC and task state FC. An independent samples *t*-test between the ASD and TD groups for each connectivity pair was run to identify group differences. P-values of all ROI-to-ROI pairs were FDR-corrected for multiple comparisons using the Benjamini-Hochberg procedure (Benjamini & Hochberg, 1995) as implemented in MATLAB. To examine group differences in overall patterns, the distributions of rest FC, task FC and reconfiguration for all ROI-to-ROI pairs (each averaged across all participants per group) were examined using non-parametric rank-sum and Kolmogorov-Smirnov tests.

### Similarity and Typicality Analyses

To examine interindividual variability of FC and of FC reconfiguration, similarity and typicality analyses were performed. Each participant’s connectivity pattern (FC matrix for 16 ROIs) was Pearson correlated with every other participant’s connectivity pattern. Mean similarity for each participant was calculated by averaging the Fisher *z*-transformed correlations between the participant and all other participants within the same group. Typicality was measured by averaging the correlations between each ASD participant and all TD participants. Group differences in similarity and typicality were calculated using permutation tests. These analyses were repeated for the ASD and TD subgroups.

### Correlations of FC Reconfiguration with Task Performance and Behavioral Measures

Pearson correlational analyses were performed between FC reconfiguration and task performance (mean accuracy and RT for SW and AW trials), language abilities (CELF-5 Word Classes subtest [CELF-5 WC]), and executive function (BRIEF-2 Global Executive Composite score [BRIEF-2 GEC]). All correlations were partially controlled for head motion during fMRI scans and age. Correlations were computed only with FC reconfiguration as we aimed to investigate how these behavioral measures are associated with change in FC during the task (interpreted as neural effort). To examine group differences in the overall pattern of correlation coefficients (FC reconfiguration with behavioral measures), the distributions in both groups were compared using non-parametric rank-sum tests.

## Results

### Task performance

The ASD group showed significantly lower accuracy for SW and AW trials, compared with the TD group, and significantly higher RT for SW trials (marginally higher for AW; Table 2). Within-group analyses further showed significantly lower accuracy for AW compared with SW and PW categories for both ASD and TD groups (Table 3). A within-group analysis of RT revealed no differences between word categories.

### Functional Connectivity and Reconfiguration

Independent samples *t*-tests between ASD and TD groups revealed no significant differences in rest FC, task FC, and FC reconfiguration for any of the ROI pairs after FDR correction (at *p*<.05; Supplemental Table S6). Non-parametric rank-sum tests were therefore used to examine group differences in the overall distributions of FC estimates (Table 4). These analyses revealed that distributions of resting state FC and task FC were significantly more positive in the ASD than the TD group, and were also more positive in both of the ASD subgroups than in the TD comparison group (Table 4; Figure 2). In addition, TP-ASD showed a more positive distribution of rest FC than LP-ASD. FC reconfiguration was greater in the full ASD sample, as well as each ASD subgroup, than in the TD comparison samples. Furthermore, reconfiguration was greater in the LP than the TP-ASD subgroup.

**Figure 2.**
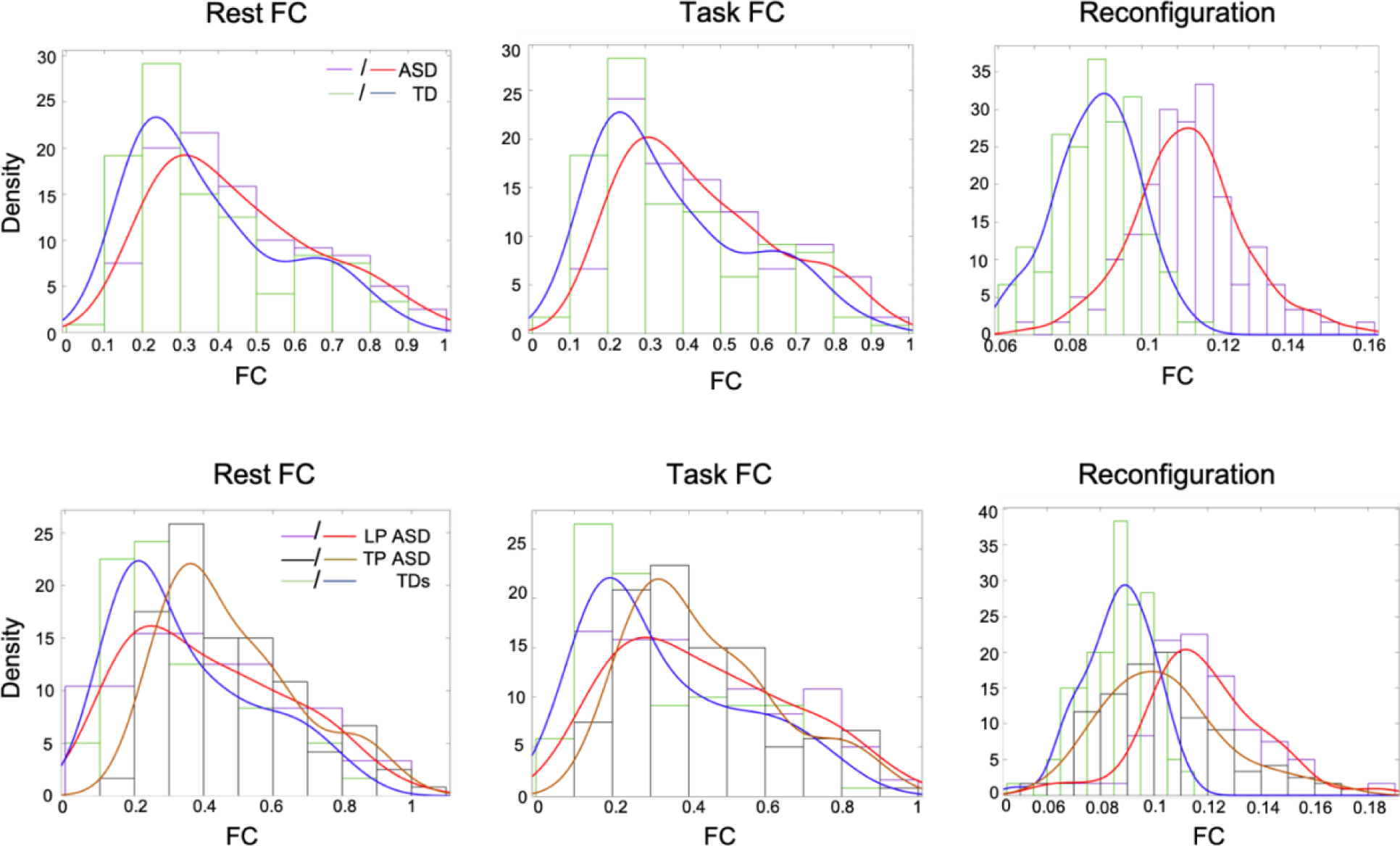
Distributions of rest FC, task FC, and reconfiguration for ASD and TD groups (top row); TP-ASD, LP-ASD, and TDs (bottom row). Density refers to the number of ROI-to-ROI pairs averaged across all participants within each group.

**Table 4.** Functional Connectivity per Sample and Non-parametric Sample Comparisons

|  |  | **TD** | **ASD** | **TD<sub>s</sub>** | **TP-ASD** | **LP-ASD** |
| --- | --- | --- | --- | --- | --- | --- |
| **Median**<br>**(SE)** | **Rest FC** | 0.30 (0.20) | 0.41 (0.21) | 0.27 (0.21) | 0.43 (0.20) | 0.38 (0.23) |
|  | **Task FC** | 0.30 (0.20) | 0.40 (0.20) | 0.29 (0.21) | 0.39 (0.19) | 0.41 (0.22) |
|  | **FC Reconfiguration** | 0.09 (0.01) | 0.11 (0.02) | 0.09 (0.01) | 0.11 (0.02) | 0.12 (0.02) |

|  |  | **ASD vs TD** | **LP-ASD vs**<br>**TD<sub>s</sub>** | **TP-ASD vs**<br>**TD<sub>s</sub>** | **LP-ASD vs**<br>**TP-ASD** |
| --- | --- | --- | --- | --- | --- |
| **z,**<br>**p -**<br>**value** | **Rest FC** | 3.2,<br>**$p = 0.002$ | 2.3,<br>*$p = 0.02$ | 5.3,<br>***$p < 0.001$ | -2.7,<br>**$p = 0.007$ |
| | **Task FC** | 3.2,<br>**$p = 0.002$ | 2.7,<br>**$p = 0.01$ | 4.1,<br>***$p < 0.001$ | -1.2,<br>$p = 0.21$ |
| | **FC Reconfiguration** | 11.1,<br>***$p < 0.001$ | 11.2,<br>***$p < 0.001$ | 8.4,<br>***$p < 0.001$ | 2.9,<br>**$p = 0.004$ |

Median value, standard error (SE) and non-parametric test results ( $z$ -statistic, $p$ -values) for all brain state FC (rest, task, reconfiguration) in each group (TD, ASD) and subgroup (TD<sub>s</sub>, TP-ASD, LP-ASD). All $p$ -values are FDR-corrected. \*\*\* $p < .001$ , \*\* $p < .01$ \* $p < .05$ .

### Similarity and Typicality of Functional Connectivity

Permutation tests demonstrated significantly reduced FC similarity in the ASD compared with the TD group for both resting state and task (Table 5). Significantly reduced similarity compared with the TD subgroup was also detected in LP-ASD for resting state only, and in TP-ASD for both resting and task state. LP-ASD and TP-ASD did not differ in similarity in any brain state FC comparison.

**Table 5.**
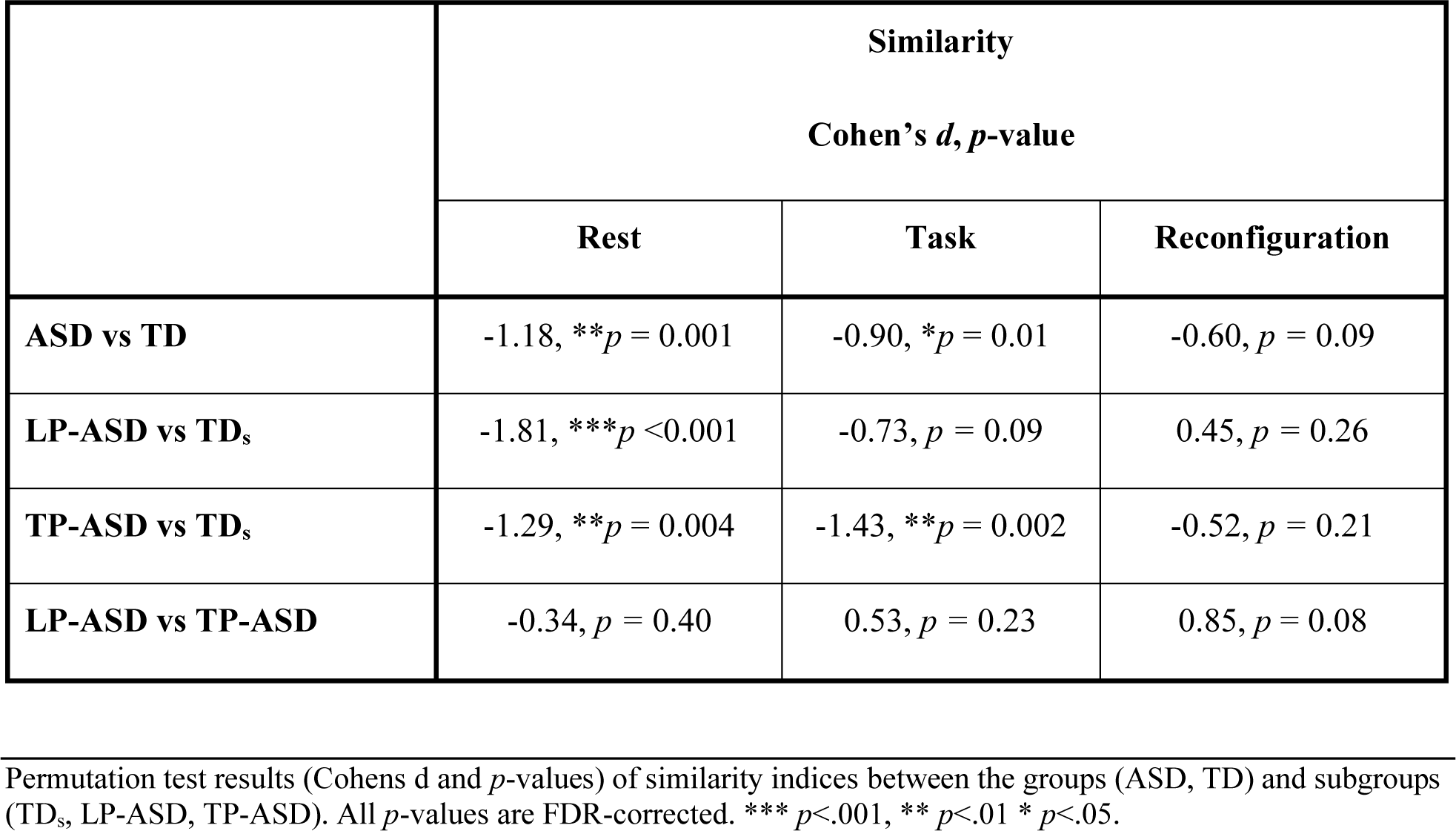
Permutation Test Results of Similarity Analyses

**Table 6.** Permutation Test Results of Typicality Analyses

|  | Typicality |  |  |
| --- | --- | --- | --- |
|  | Cohens d, *p*-value |  |  |
|  | Rest | Task | Reconfiguration |
| **ASD vs TD** | -0.45, *p* = 0.21 | -0.47, *p* = 0.21 | -0.20, *p* = 0.49 |
| **LP-ASD vs TD<sub>s</sub>** | -0.89, * *p* = 0.04 | -0.42, *p* = 0.29 | 0.18, *p* = 0.62 |
| **TP-ASD vs TD<sub>s</sub>** | -0.53, *p* = 0.21 | -0.96, * *p* = 0.03 | -0.38, *p* = 0.33 |

Permutation test results (Cohens d and *p*-values) of typicality indices between the groups (ASD, TD) and subgroups (TD<sub>s</sub>, TP-ASD, LP-ASD). All *p*-values are FDR-corrected. \* *p* < .05.

Permutation tests further revealed no significant typicality effects for rest, task, or reconfiguration (i.e., on average, FC patterns were not less similar between participants across groups than between participants within the TD group). However, when examining subgroups, reduced typicality was found in LP-ASD participants for rest FC and TP-ASD participants for task FC compared with TDs participants (Table 6).

### Correlation of Reconfiguration with Behavioral Measures

Non-parametric rank-sum tests were used to examine group differences in the relationship between FC reconfiguration and performance accuracy, RT, CELF-5 WC scores and BRIEF-2 GEC scores. The ASD group and subgroups showed more negative distributions of correlations with accuracy compared with the TD comparison groups (Table 7; Figure 3; Supplemental Figure S4). Additionally, the distributions of correlations with RT were significantly more positive in the ASD group and LP-ASD compared with the TD group and subgroup. The ASD group and both ASD subgroups further showed more negative correlations with BRIEF-GEC scores. Lastly, for correlations with CELF-5 WC, there were more positive correlations in the ASD group and TP-ASD compared with the TD group and subgroup.

**Figure 3.**
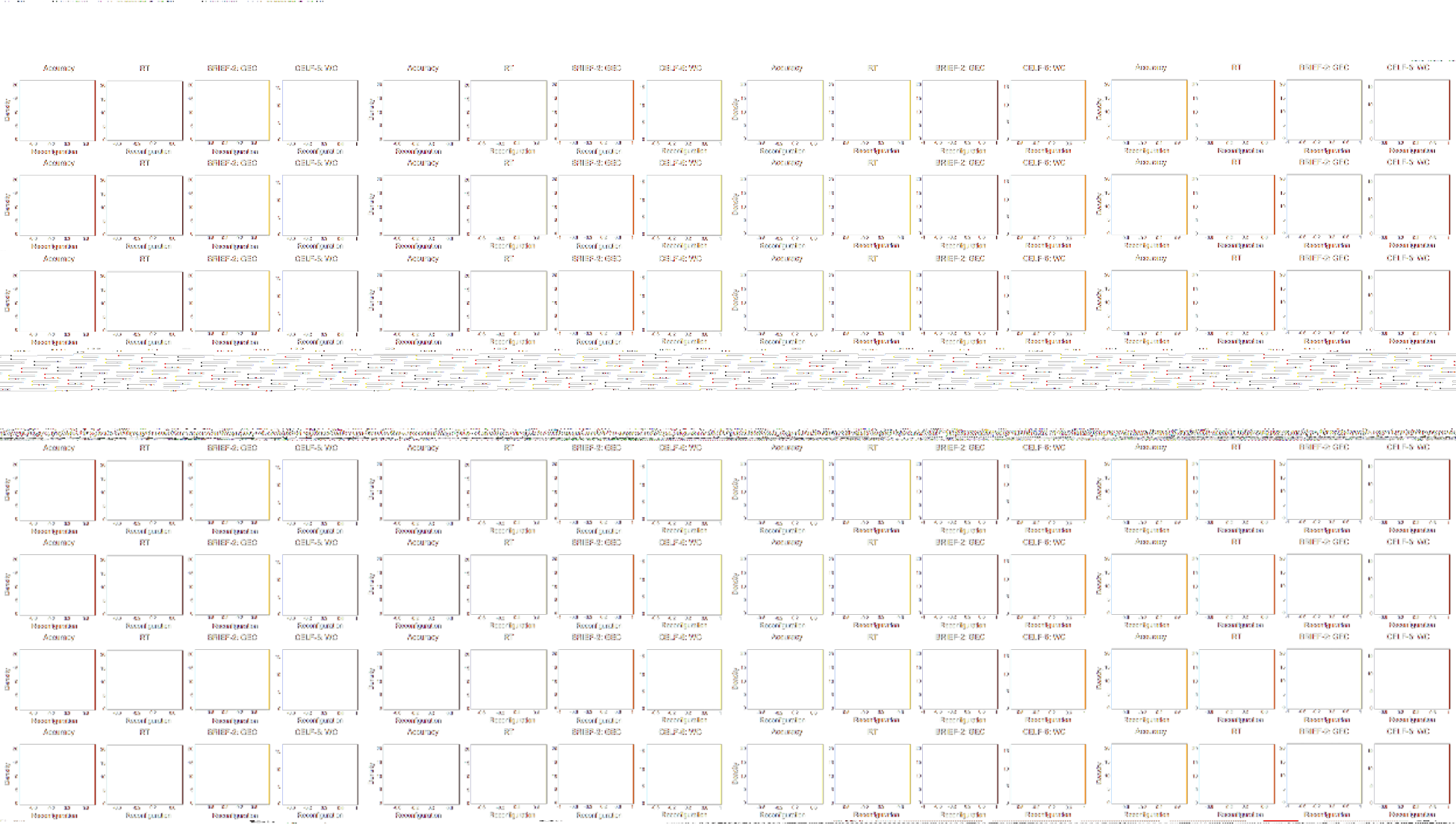
Distributions of correlation coefficients of FC reconfiguration with behavioral measures [accuracy, RT, BRIEF-2 GEC, CELF-5 WC] for ASD and TD (top row); TP-ASD, LP-ASD and TDs (bottom row).

**Table 7.** Correlations between Reconfiguration and Behavioral Measures and Non-Parametric Tests of Between-(Sub)group differences

|  | **TD** | **ASD** | **TD<sub>s</sub>** | **TP-ASD** | **LP-ASD** |
| --- | --- | --- | --- | --- | --- |
| *Correlation with:* | **Median (SE)** |  |  |  |  |
| **Accuracy** | 0.01 (0.19) | -0.06 (0.20) | 0.00 (0.23) | -0.13 (0.30) | -0.28 (0.25) |
| **RT** | 0.01 (0.23) | 0.11 (0.21) | 0.11 (0.23) | 0.13 (0.32) | 0.17 (0.26) |
| **BRIE-2 GEC** | 0.04 (0.20) | -0.16 (0.21) | 0.17 (0.28) | -0.25 (0.36) | -0.15 (0.26) |
| **CELF-5 WC** | -0.01 (0.24) | 0.11 (0.22) | 0.03 (0.27) | 0.27 (0.33) | -0.03 (0.27) |

|  | **ASD vs TD** | **LP-ASD vs TD<sub>s</sub>** | **TP-ASD vs TD<sub>s</sub>** | **LP-ASD vs TP-ASD** |
| --- | --- | --- | --- | --- |
| *Correlation with:* | ***z, p - value*** |  |  |  |
| **Accuracy** | -3.8, ***$p < 0.001$ | -7.5, ***$p < 0.001$ | -2.8, **$p = 0.008$ | -4.3, ***$p < 0.001$ |
| **RT** | 3.9, ***$p < 0.001$ | 2.1, $p = 0.04$ | 0.0, $p = 0.99$ | 1.8, $p = 0.09$ |
| **BRIEF-2 GEC** | -7.0, ***$p < 0.001$ | -7.7, ***$p < 0.001$ | -7.5, ***$p < 0.001$ | 1.8, $p = 0.09$ |
| **CELF-5 WC** | 2.9, **$p = 0.007$ | -1.6, $p = 0.11$ | 5.4, ***$p < 0.001$ | -6.5, ***$p < 0.001$ |

Median value, standard error (SE) and non-parametric test results ( $z$ -statistic, $p$ -values) for correlations of FC reconfiguration with [SW, AW] accuracy, response time (RT), BRIEF-GEC scores, and CELF-WC scores in each group (TD, ASD) and subgroup (TD<sub>s</sub>, TP-ASD, LP-ASD). All $p$ -values are FDR-corrected. \*\*\* $p < .001$ , \*\* $p < .01$ \* $p < .05$ .

## Discussion

The current investigation is among the first ASD studies to directly contrast FC during rest and task conditions. We found an overall pattern of predominant overconnectivity during both rest and task in adolescents with ASD, which is in contrast to predominant underconnectivity often reported in ASD (Di Martino et al., 2014; Duan et al., 2017; Just et al., 2012). Additionally, our ASD group showed overall greater reconfiguration than TD peers.

Dividing the ASD sample into typically-performing (TP) and low-performing (LP) subgroups shed further light on these findings: Overconnectivity was more strongly driven by the TP-ASD subgroup, while increased reconfiguration was driven by the LP-ASD subgroup. The ASD group also revealed greater heterogeneity of resting state FC and task-induced FC compared with the TD group. Reconfiguration in the TP-ASD subgroup was positively associated with the ability to understand relationships between words based on semantic class features, while no such correlation was found among TD participants. Greater reconfiguration and increased heterogeneity of FC patterns in ASD may support findings of inefficient intrinsic architecture (Dajani & Uddin, 2016; Kennedy & Courchesne, 2008), as well as recruitment of potential compensatory mechanisms (Livingston & Happé, 2017).

### Predominant Overconnectivity in ASD

The ASD group showed overall predominant overconnectivity during rest and lexical decision making – an effect that was primarily driven by the typically-performing ASD subgroup. Although underconnectivity of neural networks in ASD has long been reported (Hughes, 2007), this notion has been challenged by reports of overconnectivity (Hull et al., 2017; Picci et al., 2016), with some studies finding overconnectivity associated with greater levels of social deficits in ASD (Fishman et al., 2014; Keown et al., 2013; Supekar et al., 2013). Although overconnectivity in ASD has been interpreted as a reflection of reduced functional segregation and greater ‘cross talk’ between networks (Fishman et al., 2014, 2015; Rudie et al., 2012; Shih et al., 2011), our finding of greater overconnectivity in typically-performing than in low-performing ASD participants suggests some beneficial behavioral effects of increased FC within the language network investigated here. High levels of interregional signal correlation (i.e., strong FC) are generally driven by high amplitude events (Esfahlani et al., 2020), presumably indicating greater neural activity. This may reflect greater neural activity in the typically-performing ASD subgroup suggesting the recruitment of compensatory mechanisms for better task performance in ASD.

### Reconfiguration is Broadly Increased in ASD

Divergent results in the FC literature of ASD may be in part due to differences in methodological factors such as differences between resting and task states (Jones et al., 2010; Müller et al., 2011; Nair et al., 2014). Reconfiguration approaches these differences directly, thus opening up a complementary perspective on FC and providing added insight into neural networks in ASD. For instance, You and colleagues (2013) found atypical modulation of functional connectivity patterns from resting state to attentional brain state in children with ASD, which is consistent with other studies demonstrating atypical FC pattern changes across cognitive states in ASD (Barttfeld et al., 2012). The current study is more comprehensive as it examines FC both during rest and task, as well as FC changes between rest and task. Here, no significant group differences in reconfiguration were observed at the level of specific ROI-to-ROI connectivity. This may be due to a selection of high-functioning ASD individuals who were able to perform above 60% accuracy without extensive motion inside the scanner. Moreover, given the large number of FC comparisons and need for multiple-comparison correction (Lieberman & Cunningham, 2009), sample size may have been insufficient for detecting subtle group differences in unique ROI-to-ROI connectivities.

However, the distribution of FC reconfiguration across all ROI-to-ROI pairs (examined using rank-sum tests) revealed overall greater reconfiguration in the full ASD sample as well as in the ASD subgroups, compared to TD comparison samples. Hearne and colleagues (2017) suggested that reconfiguration increases when the system is pushed to its limits. Thus, our findings of overall stronger FC reconfiguration may indicate that ASD participants were able to perform the lexical decision task through greater neural effort. Moreover, greater FC reconfiguration in the low-performing ASD subgroup than the typically-performing ASD subgroup suggests that although the TP-ASD subgroup showed superior task performance, LP-ASD participants required greater neural change (effort) to be able to perform the task even at lower accuracy levels.

### Typical Levels of Performance in ASD may be Achieved in ‘Idiosyncratic’ Ways

Hahamy and colleagues (2015) proposed that idiosyncratic variability of functional networks may be a characteristic of the ASD brain. This is in line with some more recent studies demonstrating greater FC variability in ASD (Dickie et al., 2018; Nunes et al., 2019). In our study, FC similarity was significantly reduced in the ASD group compared to similarity within the TD group for both resting state and task. This suggests that the ASD brain may be characterized by increased interindividual variability of FC patterns, which may be considered an alternative quantitative metric of neural network abnormalities in ASD.

Further analyses of subgroups revealed greater FC variability and reduced typicality for the task condition in the typically-performing ASD subgroup, but for resting state in the low-performing ASD subgroup. This suggests that intrinsic FC architecture is atypical and idiosyncratic in low-performing ASD participants, whereas higher-performing ASD individuals may recruit idiosyncratic (potentially compensatory) mechanisms to achieve typical levels of lexical performance. Eigsti and colleagues (2016) found that children once diagnosed with ASD who no longer met diagnostic criteria and performed typically on a language comprehension task showed atypical activation in language areas. Thus, while the TD brain has robust functional networks optimized for lexical processing (Friederici, 2011), ASD individuals may recruit atypical or idiosyncratic mechanisms to achieve seemingly typical levels of task performance.

### Reconfiguration is Positively Associated with Executive Functioning but has a Complex Relation with Language Abilities in ASD

Previous behavioral studies have reported atypical performance on lexicosemantic decision and executive tasks in ASD (e.g., de Vries & Geurts, 2012; Ellis Weismer et al., 2018). The current finding of significantly lower accuracy in ASD compared to TD participants is consistent with previous studies (Dunn et al., 1999; Toichi & Kamio, 2001), suggesting that lexicosemantic processing is affected even in high-functioning individuals with autism. Overall, our ASD samples showed more positive correlations between FC reconfiguration and executive functioning, suggesting that greater reconfiguration of neural networks in ASD may be associated with better executive function, i.e., reduced impairments in regulating behavior, emotional responses, and cognitive processes (Gioia et al., 2000). As reconfiguration reflects the switching from intrinsic network connectivity to task related network connectivity, it can be presumed to require top-down control, which taps into executive abilities. This suggests that ASD adolescents with lower executive control may also have a reduced ability to reconfigure their FC. Uddin and colleagues (2015) found that lower FC reconfiguration was associated with increased severity of restricted and repetitive behaviors (RRBs) in ASD children, reflecting behavioral inflexibility, possibly indicating low executive control to limit RRBs. Greater reconfiguration in ASD may, thus, reflect greater neural flexibility associated with relatively good executive functioning.

The relation between FC reconfiguration and linguistic abilities was found to be complex. On the one hand, greater reconfiguration was associated with increased word association skills (on the CELF-5) in the full ASD sample and the typically-performing ASD subgroup. This finding suggests that some ASD individuals may achieve typical language scores through compensatory mechanisms requiring high levels of neural effort, which are however not associated with changes in intrinsic (resting state) functional network architecture. In these cases, effortful remedial strategies for achieving high levels of language processing may persist, reflected in high FC reconfiguration.

On the other hand, the distribution of correlations between FC reconfiguration and task accuracy was more negative in the ASD than the TD group. When examining the ASD subgroups, however, the shift of the distribution toward negative correlations was driven by the low-performing ASD subgroup, indicating that greater FC reconfiguration was not beneficial for task performance in this subgroup. Although the lexicosemantic system in both subgroups may have been ‘pushed to its limits’ (Hearne et al., 2017), the effect was more pronounced in the low-performing ASD subgroup. Greater neural effort in this subgroup may thus have been employed in a non-efficient way (You et al., 2013), resulting in a robustly negative correlation between reconfiguration and accuracy.

### Limitations and Future Directions

ROIs were selected based on regions showing the greatest task-related activation across ASD and TD groups, and some idiosyncratic FC patterns in ASD involving other ROIs may therefore have been missed. Moreover, useable, low-motion fMRI data could only be acquired from participants who were able to follow explicit instructions and remain still for an extended duration during rest and task scans. Our ASD sample may therefore not be representative of individuals at the lower end of the spectrum. Lastly, parent-report measures of BRIEF-2 may not fully capture measures of executive dysfunction (e.g., due to social desirability bias). Therefore, more objective behavioral measures of executive functioning abilities will be desirable in future studies.

## Conclusion

Previous studies of ASD have investigated either resting state or task state FC, with often inconsistent results. Here, we show that additional examination of FC reconfiguration, the *change* between rest and task FC, may be an informative complementary measure of the neural bases of lexical processing. In adolescents with ASD, we found atypically increased reconfiguration overall as well as greater interindividual variability. Links between reconfiguration and behavioral measures differed depending on the level of lexicosemantic task performance. Whereas reconfiguration in ASD participants with typical accuracy levels was positively associated with language skills, those performing at atypically low levels revealed a negative association between reconfiguration and task performance. Findings suggest that some individuals with ASD may recruit compensatory mechanisms to achieve typical levels of performance.

## Funding and Acknowledgements

This research was supported by the National Institutes of Health (R01-MH101173 to RAM). The funding sources had no role in study design, writing of the report, or the decision to submit the article for publication. The authors thank members of the Brain Development Imaging Laboratories, and the participating adolescents and families for their time and patience.

---

**Supplemental Figure S1.**
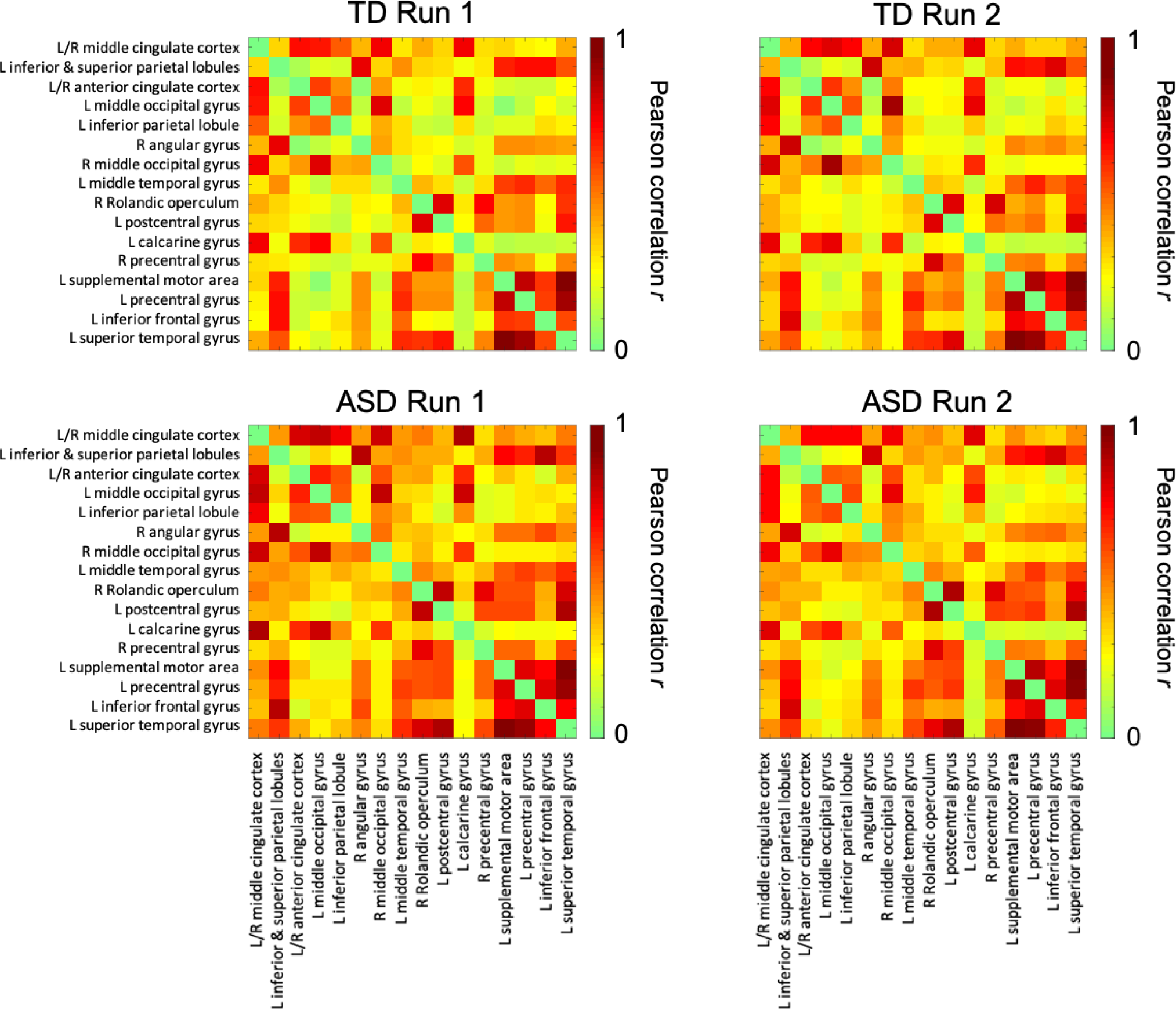
Matrix of task FC for Each Task Run

**Supplemental Figure S2.**
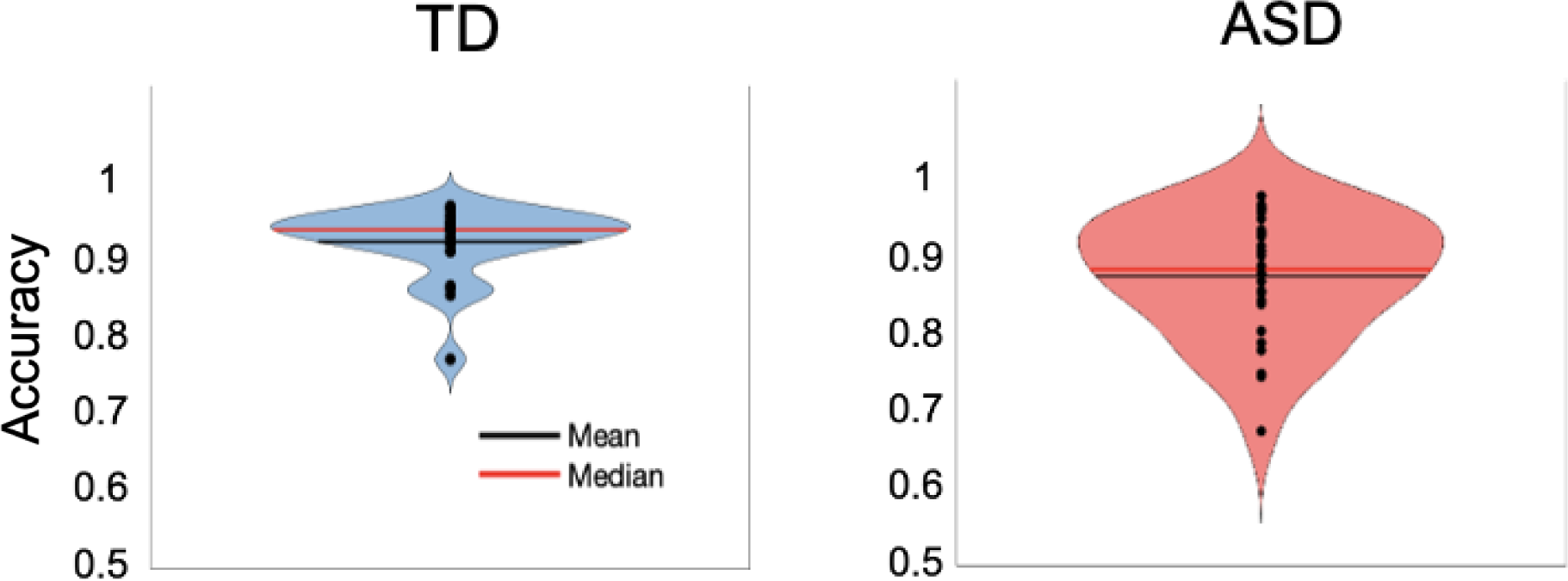
Violin Plots of Performance Accuracy in ASD and TD

**Supplemental Figure S3.**
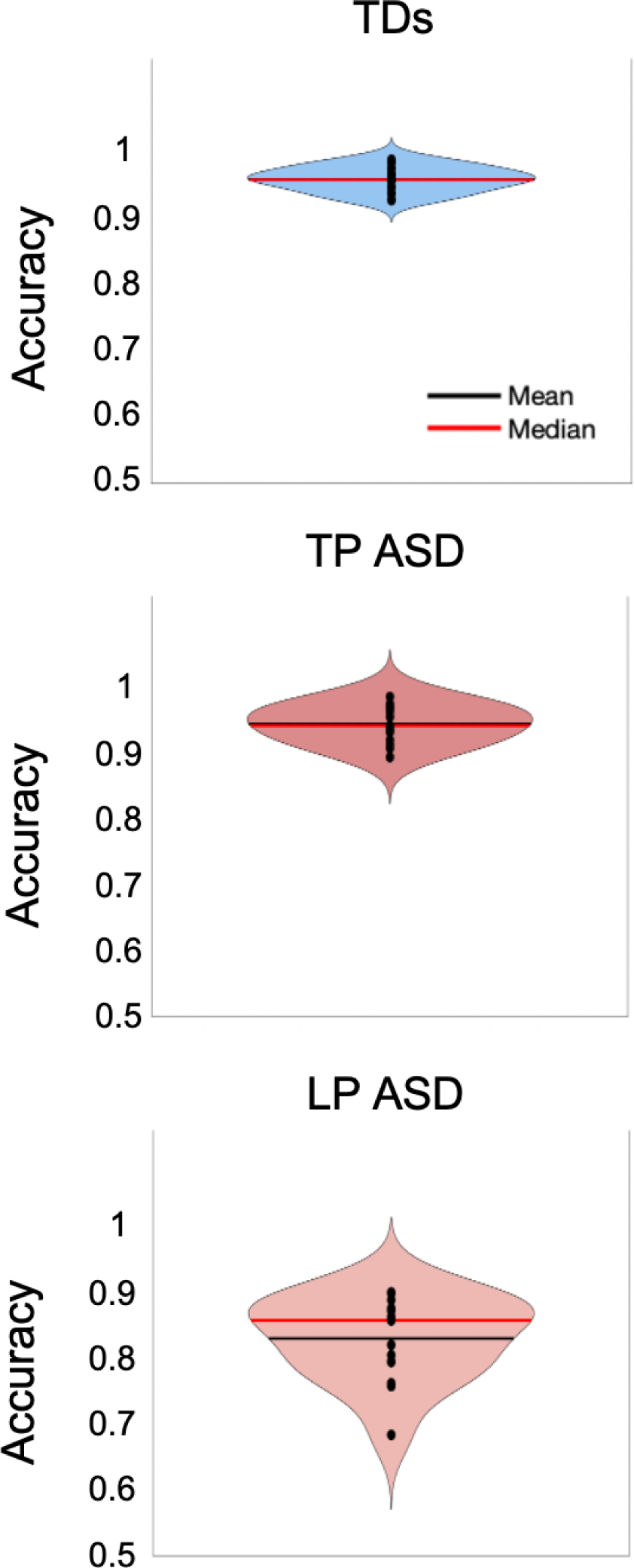
Violin Plots of Performance Accuracy in TDs, TP-ASD and LP-ASD

**Supplemental Figure S4.**
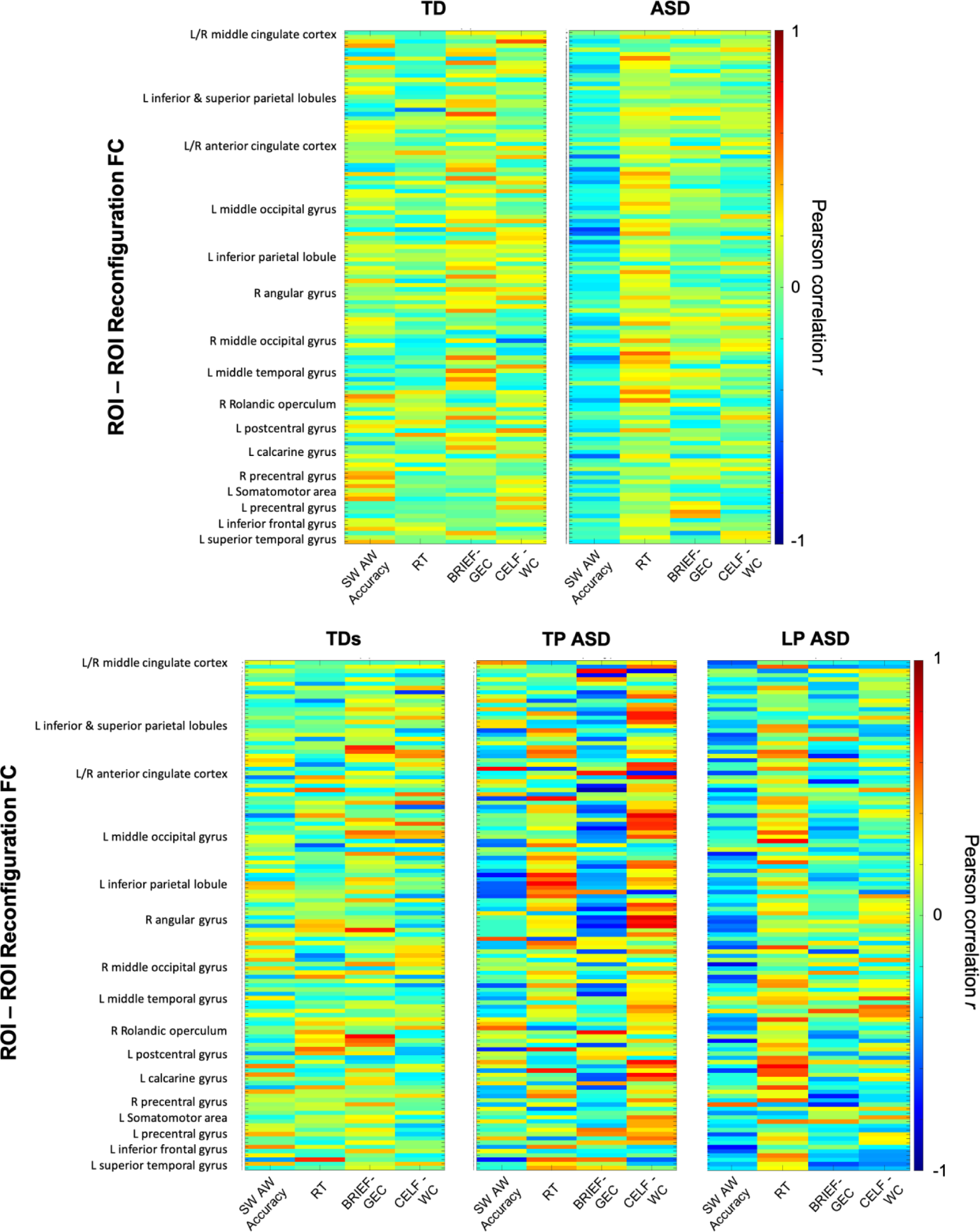
Heat maps of correlation matrices of FC reconfiguration with behavioral measures ([SW, AW] accuracy, RT, BRIEF-2 GEC scores and CELF-5 WC scores) for ASD and TD (top row); TP-ASD, LP-ASD and TDs (bottom row). Y-axis label shows the common ROI in the ROI-to-ROI pair.

**Supplemental Table S1.** Psychotropic Medication Use in the ASD Group

| Participant | Stimulants | Mood Stabilizers<sup>a</sup> | SSRI/Antidepressants | Anxiolytics/Other<sup>b</sup> | List of medications |
| --- | --- | --- | --- | --- | --- |
| 130A_V2 | + | + | + | + | Mirtazapine, Escitalopram oxalate, Aripiprazole, Methylphenidate hydrochloride |
| 420A_V2 | + |  |  | + | Methylphenidate hydrochloride, Guanfacine |
| 427A | + |  |  |  | Methylphenidate hydrochloride |
| 473A | + |  |  |  | Methylphenidate Hydrochloride |
| 490A | + | + | + | + | Oxcarbazepine, Guanfacine, Aripiprazole, Escitalopram oxalate, Alprazolam |
| 536A |  |  |  | + | Inderal |
| 542A |  | + |  |  | Aripiprazole |
| 563A |  | + | + | + | Guanfacine, Aripiprazole, Bupropion Hydrochloride |
| 577A | + | + | + |  | Aripiprazole, Lisdexamfetamine, Dextroamphetamine, Fluoxetine, Citalopram |
| 586A |  | + | + | + | Quetiapine, Sertraline, Duloxetine, Amantadine, Prazosin |
| 597A |  | + | + |  | Clomipramine, Sertaline, Bupropion, Divalproex |
| **Total** | **7** | **7** | **6** | **6** |  |

**Supplemental Table S2.** Word Category Statistics

|  |  | Frequency | Age of Acquisition (AoA) | Number of letters | Number of syllables |
| --- | --- | --- | --- | --- | --- |
| **SW** | Mean (SD) | 3.8 (0.7) | 6.8 (1.8) | 6 (2.0) | 1.9 (0.9) |
|  | Range | 2.3 – 5.6 | 3.3 -11.6 | 3 -11 | 1 -5 |
| **AW** | Mean (SD) | 3.4 (0.7) | 6.4 (1.7) | 5.9 (1.7) | 2.0 (0.7) |
|  | Range | 1.6 – 4.9 | 3.6 -10.6 | 2 -9 | 1-4 |
| **PW** | Mean (SD) | -- | -- | 6.1 (2.1) | 2.1 (1.0) |
|  | Range | -- | -- | 2-11 | 1-5 |
| **SW vs AW** | $t$ (df) =<br>$p$-values | 4.07 (238) < 0.001 | 1.43 (238) = 0.16 | 0.62 (238) = 0.53 | -0.67 (238) = 0.51 |
| **AW vs PW** | $t$ (df) =<br>$p$-values | -- | -- | -0.14 (118) = 0.89 | -1.06 (118) = 0.29 |
| **SW vs PW** | $t$ (df) =<br>$p$-values) | -- | -- | 0.09 (238) = 0.93 | -1.36 (238) = 0.17 |

**Supplemental Table S3.** Peak Coordinates of Regions of Interest

|  | **X** | **Y** | **Z** | **Voxels** |
| --- | --- | --- | --- | --- |
| **L/R middle cingulate cortex** | 9.5 | 44.5 | 30.5 | 91 |
| **L inferior & superior parietal lobules** | 24.5 | 68.5 | 33.5 | 51 |
| **L/R anterior cingulate cortex** | 6.5 | -36.5 | -11.5 | 75 |
| **L middle occipital gyrus** | 42.5 | 77.5 | 21.5 | 76 |
| **L inferior parietal lobule** | 54.5 | 59.5 | 21.5 | 84 |
| **R angular gyrus** | -32.5 | 62.5 | 39.5 | 41 |
| **R middle occipital gyrus** | -44.5 | 77.5 | 18.5 | 77 |
| **L middle temporal gyrus** | 57.5 | 32.5 | 0.5 | 64 |
| **R Rolandic operculum** | -53.5 | 17.5 | 9.5 | 66 |
| **L postcentral gyrus** | 57.5 | 20.5 | 12.5 | 53 |
| **L calcarine gyrus** | 12.5 | 59.5 | 12.5 | 54 |
| **R precentral gyrus** | -38.5 | 26.5 | 45.5 | 89 |
| **L supplemental motor area** | 6.5 | -15.5 | 36.5 | 80 |
| **L precentral gyrus** | 42.5 | 2.5 | 36.5 | 78 |
| **L inferior frontal gyrus** | 39.5 | -21.5 | 21.5 | 20 |
| **L superior temporal gyrus** | 54.5 | -12.5 | -11.5 | 88 |

**Supplemental Table S4.** Subgroup Demographics

|  | **TD<sub>s</sub> (n = 19)** |  | **LP-ASD (n =15)** |  | **TP-ASD (n = 15)** |  |
| --- | --- | --- | --- | --- | --- | --- |
| **Gender** | 6 Female |  | 4 Female |  | 5 Female |  |
| **Handedness** | 2 left |  | 0 left |  | 1 left |  |
|  | Mean (SD) | Range | Mean (SD) | Range | Mean (SD) | Range |
| **Age in years** | 15.8 (2.3) | 12.1 - 21.0 | 15.33 (2.6) | 10.0 - 20.0 | 15.67 (2.7) | 12.1 – 20.0 |
| **RMSD** | 0.06 (0.02) | 0.03 - 0.12 | 0.08 (0.02) | 0.04 - 0.11 | 0.07 (0.02) | 0.03 – 0.09 |
| **WASI-II** |  |  |  |  |  |  |
| Nonverbal IQ | 112.0 (9.9) | 90 - 128 | 106.1 (22.4) | 62 -156 | 108.1 (21.6) | 62 - 132 |
| Verbal IQ | 113.1 (12.8) | 85 - 135 | 98.4 (17.8) | 68 -124 | 110.4 (16.7) | 68 - 134 |
| Full Scale | 114.9 (12.0) | 93 - 135 | 100.3 (19.6) | 54 -136 | 111.1 (21.7) | 54 - 141 |
| **WIAT-III** | 111.2 (8.5) | 100 - 129 | 93.3 (19.0) | 59 -128 | 109.0 (18.0) | 59 - 133 |
| **ADOS-2 Total<sup>†</sup>** | -- | -- | 10.9 (3.7) | 6 - 20 | 10.6 (3.8) | 2 - 16 |
| **BRIEF-2 GEC** | 45.0 (7.7) | 36 - 62 | 66.5 (5.8) | 56 - 74 | 65.9 (9.6) | 49 - 84 |
| **CELF-5 WC** | 36.2 (2.2) | 32 - 39 | 34.0 (3.8) | 27 -39 | 35.9 (4.4) | 27 - 40 |
| **Psychotropic medication use** | -- |  | 6 reported |  | 5 reported |  |

**Supplemental Table S5.** Subgroup Demographic Statistics

|  | LP-ASD vs TD<sub>s</sub> | TP-ASD vs TD<sub>s</sub> | LP-ASD vs TP-ASD |
| --- | --- | --- | --- |
|  | *t* (df), *p* - value |  |  |
| **Age in years** | 0.4 (32), *p* = 0.69 | 0.8 (32), *p* = 0.41 | -0.4 (28), *p* = 0.73 |
| **RMSD** | 2.2 (32), *p* = 0.03 | 1.3 (32), *p* = 0.21 | -1.2 (28), *p* = 0.25 |
| **WASI-II** |  |  |  |
| Nonverbal IQ | -1.0 (32), *p* = 0.31 | -0.7 (32), *p* = 0.49 | -0.2 (28), *p* = 0.81 |
| Verbal IQ | -2.8 (32), ** *p* = 0.01 | -0.5 (32), *p* = 0.60 | -1.9 (28), *p* = 0.07 |
| Full Scale | -2.7 (32), * *p* = 0.01 | -0.7 (32), *p* = 0.52 | -1.4 (28), *p* = 0.16 |
| **WIAT-III** | -3.7 (32), ** *p* = 0.001 | -0.5 (32), *p* = 0.64 | -2.3 (28), * *p* = 0.03 |
| **ADOS-2 Total<sup>†</sup>** | -- | -- | 0.2 (28), *p* = 0.81 |
| **BRIEF-2 GEC** | 8.7 (30), *** *p* < 0.001 | 6.3 (26), *** *p* < 0.001 | 0.2 (22), *p* = 0.85 |
| **CELF-5 WC** | -2.2 (32), * *p* = 0.04 | -0.3 (32), *p* = 0.78 | -1.3 (28), *p* = 0.21 |

\*\*\* *p* < .001, \*\* *p* < .01 \* *p* < .05.Supplemental Table S6. FDR-corrected *p*- values for Group Differences in Functional Connectivity

**Supplemental Table S6.** FDR-corrected *p-* values for Group Differences in Functional Connectivity

| Connectivity (ROI pairing) | t- statistic (df) = *p* - values |  |  |
| --- | --- | --- | --- |
|  | Rest | Task | Reconfiguration |
| L/R middle cingulate cortex-L inferior & superior parietal lobules | 0.52(51) = 0.28 | 0.98(51) = 0.33 | 0.74(51) = 0.46 |
| L/R middle cingulate cortex-L/R anterior cingulate cortex | 0.92(51) = 0.46 | 1.19(51) = 0.24 | 1.43(51) = 0.16 |
| L/R middle cingulate cortex-L middle occipital gyrus | 0.39(51) = 0.25 | 1.32(51) = 0.19 | 0.94(51) = 0.35 |
| L/R middle cingulate cortex-L inferior parietal lobule | 0.96(51) = 0.97 | 2.01(51) = 0.05 | 1.13(51) = 0.26 |
| L/R middle cingulate cortex-R angular gyrus | 0.87(51) = 0.72 | 0.89(51) = 0.38 | 0.67(51) = 0.5 |
| L/R middle cingulate cortex-R angular gyrus | 0.35(51) = 0.12 | 0.64(51) = 0.52 | 1.05(51) = 0.3 |
| L/R middle cingulate cortex-L middle temporal gyrus | 2.28(51) = 0.56 | 2.31(51) = 0.03 | -0.18(51) = 0.86 |
| L/R middle cingulate cortex-R Rolandic operculum | 1.47(51) = 0.13 | 1.51(51) = 0.14 | 0.35(51) = 0.73 |
| L/R middle cingulate cortex-L postcentral gyrus | 0.1(51) = 0.57 | 0.41(51) = 0.68 | 1.53(51) = 0.13 |
| L/R middle cingulate cortex-L calcarine gyrus | 1.69(51) = 0.06 | 1.96(51) = 0.06 | 0.36(51) = 0.72 |
| L/R middle cingulate cortex-R precentral gyrus | 0.14(51) = 0.56 | 0.13(51) = 0.89 | 2.02(51) = 0.05 |
| L/R middle cingulate cortex-L SMA, L middle cingulate cortex | 1.23(51) = 0.55 | 1.44(51) = 0.15 | -0.01(51) = 0.99 |
| L/R middle cingulate cortex-L precentral gyrus | 0.97(51) = 0.92 | 1.33(51) = 0.19 | 0.29(51) = 0.77 |
| L/R middle cingulate cortex-L inferior frontal gyrus | 0.32(51) = 0.31 | 0.82(51) = 0.42 | 0.38(51) = 0.71 |
| L/R middle cingulate cortex-L superior temporal gyrus | 0.89(51) = 0.74 | 1.06(51) = 0.29 | 0.73(51) = 0.47 |
| L inferior & superior parietal lobules-L/R anterior cingulate cortex | 0.38(51) = 0.61 | 0.69(51) = 0.49 | 1.46(51) = 0.15 |
| L inferior & superior parietal lobules-L middle occipital gyrus | 0.51(51) = 0.8 | 1.11(51) = 0.27 | 2.46(51) = 0.02 |
| L inferior & superior parietal lobules-L inferior parietal lobule | 1.16(51) = 0.42 | 0.91(51) = 0.37 | 1.39(51) = 0.17 |
| L inferior & superior parietal lobules-R angular gyrus | 0.83(51) = 0.99 | 1.2(51) = 0.24 | 1.89(51) = 0.06 |
| L inferior & superior parietal lobules-R angular gyrus | -0.36(51) = 0.83 | 0.58(51) = 0.56 | 1.99(51) = 0.05 |
| L inferior & superior parietal lobules-L middle temporal gyrus | 1.15(51) = 0.99 | 0.54(51) = 0.59 | 0.74(51) = 0.46 |
| L inferior & superior parietal lobules-R Rolandic operculum | 1.32(51) = 0.89 | 1.19(51) = 0.24 | 1.78(51) = 0.08 |
| L inferior & superior parietal lobules-L postcentral gyrus | 1.56(51) = 0.48 | 1.52(51) = 0.13 | 1.44(51) = 0.16 |
| L inferior & superior parietal lobules-L calcarine gyrus | 0.5(51) = 0.45 | 0.53(51) = 0.6 | 1(51) = 0.32 |
| L inferior & superior parietal lobules-R precentral gyrus | 0.93(51) = 0.97 | 1.41(51) = 0.16 | 1.04(51) = 0.3 |
| L inferior & superior parietal lobules-L SMA, L middle cingulate cortex | 0.6(51) = 0.79 | 0.56(51) = 0.57 | 2.75(51) = 0.01 |
| L inferior & superior parietal lobules-L precentral gyrus | 1.19(51) = 0.71 | 0.42(51) = 0.67 | 1.77(51) = 0.08 |
| L inferior & superior parietal lobules-L inferior frontal gyrus | 1.04(51) = 0.83 | 1.45(51) = 0.15 | 2.72(51) = 0.01 |
| L inferior & superior parietal lobules-L superior temporal gyrus | 1.43(51) = 0.81 | 1.05(51) = 0.3 | 2.22(51) = 0.03 |
| L/R anterior cingulate cortex-L middle occipital gyrus | -0.2(51) = 0.51 | 0.08(51) = 0.94 | 2.03(51) = 0.05 |
| L/R anterior cingulate cortex-L inferior parietal lobule | 1.7(51) = 0.33 | 1.94(51) = 0.06 | 1.86(51) = 0.07 |
| L/R anterior cingulate cortex-R angular gyrus | 1.57(51) = 0.92 | 1.76(51) = 0.08 | 1.69(51) = 0.1 |
| L/R anterior cingulate cortex-R angular gyrus | 2.25(51) = 0.41 | 2.3(51) = 0.03 | 0.68(51) = 0.5 |
| L/R anterior cingulate cortex-L middle temporal gyrus | 1.25(51) = 0.53 | 1.99(51) = 0.05 | 0.78(51) = 0.44 |
| L/R anterior cingulate cortex-R Rolandic operculum | 2.33(51) = 0.1 | 2.62(51) = 0.01 | 0.79(51) = 0.43 |
| L/R anterior cingulate cortex-L postcentral gyrus | 0.5(51) = 0.17 | 0.69(51) = 0.49 | 1.41(51) = 0.17 |
| L/R anterior cingulate cortex-L calcarine gyrus | -0.11(51) = 0.9 | -0.11(51) = 0.92 | 1.96(51) = 0.06 |
| L/R anterior cingulate cortex-R precentral gyrus | 1.14(51) = 0.13 | 0.86(51) = 0.39 | 1.74(51) = 0.09 |
| L/R anterior cingulate cortex-L SMA, L middle cingulate cortex | 1.05(51) = 0.25 | 1.84(51) = 0.07 | 1.33(51) = 0.19 |
| L/R anterior cingulate cortex-L precentral gyrus | 0.67(51) = 0.82 | 1.35(51) = 0.18 | 1.61(51) = 0.11 |
| L/R anterior cingulate cortex-L inferior frontal gyrus | 0.62(51) = 0.46 | 0.49(51) = 0.63 | 1.02(51) = 0.31 |
| L/R anterior cingulate cortex-L superior temporal gyrus | 0.86(51) = 0.17 | 1.49(51) = 0.14 | 1.01(51) = 0.32 |
| L middle occipital gyrus-L inferior parietal lobule | 0.96(51) = 0.88 | 0.49(51) = 0.63 | 0.4(51) = 0.69 |
| L middle occipital gyrus-R angular gyrus | 1.24(51) = 0.85 | 1.55(51) = 0.13 | 1.34(51) = 0.19 |
| L middle occipital gyrus-R angular gyrus | -0.61(51) = 0.11 | 0.15(51) = 0.88 | 0.32(51) = 0.75 |
| L middle occipital gyrus-L middle temporal gyrus | 1.83(51) = 0.25 | 2.51(51) = 0.02 | 0.62(51) = 0.54 |
| L middle occipital gyrus-R Rolandic operculum | 0.76(51) = 0.76 | 1.17(51) = 0.25 | 1.13(51) = 0.27 |
| L middle occipital gyrus-L postcentral gyrus | -0.33(51) = 0.46 | 0.08(51) = 0.94 | 1.61(51) = 0.11 |
| L middle occipital gyrus-L calcarine gyrus | 0.02(51) = 0.92 | 0.73(51) = 0.47 | -0.77(51) = 0.44 |
| L middle occipital gyrus-R precentral gyrus | 0.28(51) = 0.89 | 0.35(51) = 0.72 | 1.81(51) = 0.08 |
| L middle occipital gyrus-L SMA, L middle cingulate cortex | 0.63(51) = 0.74 | 1.47(51) = 0.15 | 0.82(51) = 0.42 |
| L middle occipital gyrus-L precentral gyrus | 1.14(51) = 0.2 | 1.87(51) = 0.07 | 1.83(51) = 0.07 |
| L middle occipital gyrus-L inferior frontal gyrus | 0.19(51) = 0.67 | 0.59(51) = 0.56 | -0.81(51) = 0.42 |
| L middle occipital gyrus-L superior temporal gyrus | 0.58(51) = 0.59 | 1.03(51) = 0.31 | 1.97(51) = 0.05 |
| L inferior parietal lobule-R angular gyrus | 1.34(51) = 0.87 | 1.04(51) = 0.31 | 1.81(51) = 0.08 |
| L inferior parietal lobule-R angular gyrus | 0.81(51) = 0.99 | 1.04(51) = 0.3 | 2.58(51) = 0.01 |
| L inferior parietal lobule-L middle temporal gyrus | 1.46(51) = 0.5 | 1.45(51) = 0.15 | 0.69(51) = 0.5 |
| L inferior parietal lobule-R Rolandic operculum | 1.09(51) = 0.57 | 1.46(51) = 0.15 | 1.06(51) = 0.29 |
| L inferior parietal lobule-L postcentral gyrus | 0.31(51) = 0.69 | 0.46(51) = 0.65 | 1.57(51) = 0.12 |
| L inferior parietal lobule-L calcarine gyrus | 1.59(51) = 0.94 | 2.17(51) = 0.03 | -0.74(51) = 0.46 |
| L inferior parietal lobule-R precentral gyrus | 0.46(51) = 0.87 | 0.68(51) = 0.5 | -0.14(51) = 0.89 |
| L inferior parietal lobule-L SMA, L middle cingulate cortex | 0.31(51) = 0.82 | 0.29(51) = 0.77 | 0.99(51) = 0.33 |
| L inferior parietal lobule-L precentral gyrus | 1.17(51) = 0.63 | 1.28(51) = 0.21 | 1.76(51) = 0.08 |
| L inferior parietal lobule-L inferior frontal gyrus | -0.01(51) = 0.55 | -0.29(51) = 0.77 | 1.02(51) = 0.31 |
| L inferior parietal lobule-L superior temporal gyrus | 0.68(51) = 0.75 | 0.89(51) = 0.38 | 1.08(51) = 0.28 |
| R angular gyrus-R angular gyrus | 1.09(51) = 0.89 | 1.22(51) = 0.23 | 2.57(51) = 0.01 |
| R angular gyrus-L middle temporal gyrus | 1.84(51) = 0.55 | 1.27(51) = 0.21 | 1.83(51) = 0.07 |
| R angular gyrus-R Rolandic operculum | 1.32(51) = 0.79 | 1.3(51) = 0.2 | 2.13(51) = 0.04 |
| R angular gyrus-L postcentral gyrus | 1.27(51) = 0.95 | 1.13(51) = 0.26 | 1.08(51) = 0.29 |
| R angular gyrus-L calcarine gyrus | 0.53(51) = 0.46 | 0.92(51) = 0.36 | 1.2(51) = 0.24 |
| R angular gyrus-R precentral gyrus | 0.77(51) = 0.8 | 1.04(51) = 0.31 | 1.35(51) = 0.18 |
| R angular gyrus-L SMA, L middle cingulate cortex | 0.63(51) = 0.3 | 0.67(51) = 0.5 | 1.81(51) = 0.08 |
| R angular gyrus-L precentral gyrus | 1.62(51) = 0.34 | 1.24(51) = 0.22 | 1.7(51) = 0.1 |
| R angular gyrus-L inferior frontal gyrus | 1.12(51) = 0.96 | 1.53(51) = 0.13 | 0.97(51) = 0.34 |
| R angular gyrus-L superior temporal gyrus | 0.93(51) = 0.79 | 0.79(51) = 0.43 | 2.41(51) = 0.02 |
| R angular gyrus-L middle temporal gyrus | 3(51) = 0.34 | 2.92(51) = 0.01 | 1.07(51) = 0.29 |
| R angular gyrus-R Rolandic operculum | 1.65(51) = 0.58 | 1.9(51) = 0.06 | 2.63(51) = 0.01 |
| R angular gyrus-L postcentral gyrus | 0.63(51) = 0.47 | 0.9(51) = 0.37 | 2.71(51) = 0.01 |
| R angular gyrus-L calcarine gyrus | -0.43(51) = 0.45 | 0.15(51) = 0.88 | -0.45(51) = 0.65 |
| R angular gyrus-R precentral gyrus | 0.47(51) = 0.7 | 0.47(51) = 0.64 | 2.7(51) = 0.01 |
| R angular gyrus-L SMA, L middle cingulate cortex | 1.06(51) = 0.94 | 1.39(51) = 0.17 | 1.33(51) = 0.19 |
| R angular gyrus-L precentral gyrus | 1.07(51) = 0.88 | 1.16(51) = 0.25 | 3.09(51) = 0 |
| R angular gyrus-L inferior frontal gyrus | 0.7(51) = 0.82 | 0.96(51) = 0.34 | 2.13(51) = 0.04 |
| R angular gyrus-L superior temporal gyrus | 1.26(51) = 0.63 | 1.23(51) = 0.22 | 2.17(51) = 0.03 |
| L middle temporal gyrus-R Rolandic operculum | 1.83(51) = 0.66 | 1.73(51) = 0.09 | 2.23(51) = 0.03 |
| L middle temporal gyrus-L postcentral gyrus | 1.39(51) = 0.82 | 1.52(51) = 0.14 | 0.37(51) = 0.71 |
| L middle temporal gyrus-L calcarine gyrus | 2.04(51) = 0.31 | 2.34(51) = 0.02 | -0.53(51) = 0.6 |
| L middle temporal gyrus-R precentral gyrus | 0.83(51) = 0.69 | 1.09(51) = 0.28 | 0.36(51) = 0.72 |
| L middle temporal gyrus-L SMA, L middle cingulate cortex | 0.09(51) = 0.75 | -0.42(51) = 0.68 | 0.09(51) = 0.93 |
| L middle temporal gyrus-L precentral gyrus | 0.26(51) = 0.44 | -0.23(51) = 0.82 | 1.87(51) = 0.07 |
| L middle temporal gyrus-L inferior frontal gyrus | 0.16(51) = 0.52 | 0.06(51) = 0.95 | 0.52(51) = 0.61 |
| L middle temporal gyrus-L superior temporal gyrus | 0.31(51) = 0.54 | 0.05(51) = 0.96 | 0.42(51) = 0.68 |
| R Rolandic operculum-L postcentral gyrus | 2(51) = 0.56 | 1.38(51) = 0.17 | 1.19(51) = 0.24 |
| R Rolandic operculum-L calcarine gyrus | 1.04(51) = 0.06 | 1(51) = 0.32 | 0.71(51) = 0.48 |
| R Rolandic operculum-R precentral gyrus | 0.71(51) = 0.84 | 0.58(51) = 0.57 | 1.37(51) = 0.18 |
| R Rolandic operculum-L SMA, L middle cingulate cortex | 2.41(51) = 0.29 | 2.25(51) = 0.03 | 0.02(51) = 0.99 |
| R Rolandic operculum-L precentral gyrus | 1.5(51) = 0.66 | 1.48(51) = 0.14 | 3.37(51) = 0 |
| R Rolandic operculum-L inferior frontal gyrus | 1.72(51) = 0.94 | 1.97(51) = 0.05 | 1.21(51) = 0.23 |
| R Rolandic operculum-L superior temporal gyrus | 2.08(51) = 0.39 | 2.08(51) = 0.04 | 1.27(51) = 0.21 |
| L postcentral gyrus-L calcarine gyrus | -0.22(51) = 0.31 | -0.53(51) = 0.6 | 1.18(51) = 0.25 |
| L postcentral gyrus-R precentral gyrus | 2.1(51) = 0.83 | 1.48(51) = 0.14 | 0.73(51) = 0.47 |
| L postcentral gyrus-L SMA, L middle cingulate cortex | 1.97(51) = 0.44 | 2.11(51) = 0.04 | 0.27(51) = 0.79 |
| L postcentral gyrus-L precentral gyrus | 1.8(51) = 0.94 | 1.64(51) = 0.11 | 2.06(51) = 0.04 |
| L postcentral gyrus-L inferior frontal gyrus | 1.67(51) = 0.46 | 1.92(51) = 0.06 | 0.89(51) = 0.38 |
| L postcentral gyrus-L superior temporal gyrus | 2.04(51) = 0.53 | 2.29(51) = 0.03 | -0.02(51) = 0.99 |
| L calcarine gyrus-R precentral gyrus | 0.11(51) = 0.34 | -0.32(51) = 0.75 | -0.1(51) = 0.92 |
| L calcarine gyrus-L SMA, L middle cingulate cortex | 1.09(51) = 0.41 | 1.46(51) = 0.15 | -0.06(51) = 0.95 |
| L calcarine gyrus-L precentral gyrus | 0.38(51) = 0.59 | 0.98(51) = 0.33 | 1.49(51) = 0.14 |
| L calcarine gyrus-L inferior frontal gyrus | 0.06(51) = 0.19 | 0.78(51) = 0.44 | 1.31(51) = 0.2 |
| L calcarine gyrus-L superior temporal gyrus | 1(51) = 0.32 | 1.18(51) = 0.24 | 0.94(51) = 0.35 |
| R precentral gyrus -L SMA, L middle cingulate cortex | 1.09(51) = 0.58 | 1.15(51) = 0.26 | 0.52(51) = 0.6 |
| R precentral gyrus -L precentral gyrus | 0.53(51) = 0.16 | 0.99(51) = 0.33 | 0.5(51) = 0.62 |
| R precentral gyrus -L inferior frontal gyrus | 1.35(51) = 0.89 | 1.52(51) = 0.13 | 0.88(51) = 0.38 |
| R precentral gyrus -L superior temporal gyrus | 1.6(51) = 0.7 | 1.72(51) = 0.09 | 0.66(51) = 0.51 |
| L SMA, L middle cingulate cortex-L precentral gyrus | 0.93(51) = 0.3 | -0.07(51) = 0.94 | 1.29(51) = 0.2 |
| L SMA, L middle cingulate cortex-L inferior frontal gyrus | 0.45(51) = 0.83 | 0.65(51) = 0.52 | 2.67(51) = 0.01 |
| L SMA, L middle cingulate cortex-L superior temporal gyrus | 1.36(51) = 0.87 | 0.59(51) = 0.56 | -0.27(51) = 0.79 |
| L precentral gyrus-L inferior frontal gyrus | 1.82(51) = 0.81 | 1.94(51) = 0.06 | 0.28(51) = 0.78 |
| L precentral gyrus-L superior temporal gyrus | 1.32(51) = 0.92 | 0.67(51) = 0.51 | 1.2(51) = 0.23 |
| L inferior frontal gyrus-L superior temporal gyrus | 0.95(51) = 0.91 | 1.26(51) = 0.21 | 1.27(51) = 0.21 |

All p-values are uncorrected.

## Notes

### Competing Interest Statement

The authors have declared no competing interest.

